# A method to estimate effective population size from linkage disequilibrium when generations overlap

**DOI:** 10.1101/2021.02.17.431658

**Authors:** Luis Alberto García Cortés, Frédéric Austerlitz, M. Ángeles R. de Cara

## Abstract

Effective population size (*N*_*e*_) is a key parameter in evolutionary and conservation studies. It represents the number of individuals of an ideal panmictic population that would have the same genetic drift as the observed population and can be used to establish management programmes. Several methods have been developed to estimate this parameter. Currently, for studies with one sample in time, the simplest methods are based on linkage disequilibrium. These methods rely on simple models, and biases have been shown when populations deviate from the assumptions made in those models. This occurs in particular when populations are age-structured or have overlapping generations. Recently, several methods have been developed to correct such biases. Here, we develop analytical equations to predict linkage disequilibrium within age groups, and use such results to infer cohort size from samples of newborn individuals. We can in turn use these equations to estimate *N*_*e*_ accurately for a variety of species. Furthermore, using publicly available data, we apply our method to the white-crowned sparrow (*Zonotrichia leucophrys*).

## Introduction

Linkage disequilibrium is the association between alleles at different loci within or between chromosomes. It depends on allelic frequencies, recombination, mutation and effective population size (*N*_*e*_, Hill & Robertson 1968). Thus, it can be used to make inferences about effective size, a crucial parameter in evolutionary and conservation studies. It is commonly considered that populations with *N*_*e*_ smaller than 50 are endangered in the short-term and that conservation programmes are often necessary for such populations (Franklin 1980). Most analytical predictions on linkage disequilibrium (LD) have been obtained in populations with non-overlapping generations, and such simple models have allowed a whole body of research aiming at estimating *N*_*e*_ from LD, starting with the pioneering work of Sved (1971).

Sved followed the work of Hill & Robertson (1968), which showed how alleles at different positions in the genome tend to become associated in finite populations. He studied LD between two biallelic loci and established that it is equivalent to studying identity by descent (IBD). By analysing the temporal dynamics of the allelic frequencies at the two loci ignoring mutation, he derived an analytical formula establishing the relationship between *N*_*e*_ and LD. In later work, Sved & Feldman (1973) observed that IBD and inbreeding can be calculated either using correlations between frequencies or using probability methods. This led them to take into account sampling with replacement, which resulted in a slight modification of Sved’s (1971) formula. Weir & Hill (1980) and Hill (1981) also developed methods to infer *N*_*e*_ from LD measures, taking into account sample sizes and mating systems.

This body of work was not much used for a long time on real data, with a few exceptions in *Drosophila* and human data (Loukas *et al* 1979; Malpica & Briscoe 1982; Piazza *et al* 1985). With the advent of dense genotyping data, in particular in humans and livestock, and the work of Hayes *et al* (2003) who developed methods for inferring current and ancestral *N*_*e*_, such estimates have become common to the point where they constitute the prevailing method nowadays. In parallel to theoretical works on the relationship between *N*_*e*_ and LD, there has been a growing body of research showing the effects of population and age structure on estimates of *N*_*e*_. This started with the work of Felsenstein (1971) who derived equations for *N*_*e*_ given a known age structure and Hill (1972) showed that *N*_*e*_ in such populations is equal to that of a population with non-overlapping generations with the same variance in lifetime family size. Orive (1993) used a coalescent approach to generalise to populations with overlapping generations and sexual or asexual reproduction. These works have mostly been used to calculate *N*_*e*_ in populations once their age structure has been determined, without making use of information on linkage disequilibrium.

Only recently methods using both LD and data on age structure have been developed to improve estimates of *N*_*e*_ (Robinson & Moyer 2013; Waples *et al* 2014). These works have shown that using only LD to infer *N*_*e*_ in populations with overlapping generations can result in biased estimates, and Waples *et al* (2014) introduced heuristic modifications to correct such biases.

Here we develop analytical methods to account for age structure in the estimates of *N*_*e*_ from linkage disequilibrium. We start by deriving equations for linkage disequilibrium measured via *r*^2^ (Hill & Robertson 1968) in a scenario consisting in a population with several cohorts of varying fertility and survival rates. We use the equivalence between *r*^2^ and identity by descent to relate these genetic measures to *N*_*e*_. By inverting the system of equations derived, we can then estimate the cohort size and the effective population size from genetic data. Furthermore, we take advantage of our previous research (Garcia-Cortes *et al* 2019), to show the importance of dealing appropiately with frequencies or probabilities, and how they lead to very different results. Finally, we apply our method to publicly available data from the white-crowned sparrow (*Zonotrichia leucophrys*, Lipshutz *et al* 2017).

## Methods and material

Throughout the paper, we consider populations of *N* individuals who are diploid and, therefore, correspond to populations 2*N* gametes. Sved’s (1971) method defines *Q* as the probability of identity by descent (IBD) of two gametes at locus B given that locus A is also IBD. This probability *Q* is also called the joint homozygosity and represents the probability of no crossing over between loci A and B since the common ancestor. Sved (1971) showed that *E* (*r*^2^) = *Q*, where *r*^2^ = *D*^2^/ (*p*_*A*_ *p*_*a*_ *p*_*B*_ *p*_*b*_) is a measure of linkage disequilibrium between loci A and B, and derived the following probability of joint homozygosity in the next generation, *Q*^′^, as

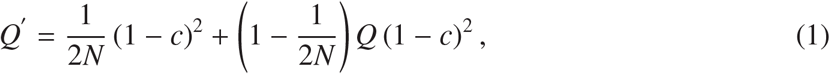

where *c* is the recombination fraction between loci *A* and *B* and *N* is the effective population size. This size is defined in Kimura & Crow (1963), as the size of an idealised population with the same inbreeding or random genetic drift as the population under consideration. The recombination fraction is obtained as *c* = 0.5 (1 − *e*^−2*d*^)where *d* is the map distance (Haldane 1919).

This equation was later revised by Sved & Feldman (1973) as

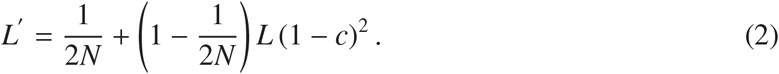

where *L* is the probability of identity by descent between loci *A* and *B*, also called linked identity by descent (Sved 2019). In the original paper where this modification was added (Sved & Feldman 1973), the rationale was that the probability of IBD for two loci follows the same logic as the probability of IBD for one locus 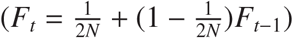. Sved (1971) had derived eq. 1 thinking of frequencies rather than probabilities.

Thus, while *Q* and *L* are similar in concept, one of them is not appropiate (Sved 2019). *Q* is the probability of joint IBD without replacement, while *L* is the probability of joint IBD, but allowing for replacement. Earlier work has shown that *L*, as derived in Sved & Feldman (1973), provides better estimations of *E* [*r*^2^] (Garcia-Cortes *et al* 2019).

Equations (1) and (2) were developed for populations with non-overlapping generations. Nevertheless, many species have overlapping generations and this cannot be ignored when evaluating the intensity of genetic drift (Felsenstein 1971), or calculating the equilibrium between genetic drift and recombination. Most importantly, researchers have used estimates of *N*_*e*_ based on these equations without much caution about potential pitfalls (Robinson & Moyer 2013; Waples *et al* 2014).

In populations with age structure, following the notation of Felsenstein (1971), the new cohort of *N*_0_ newborns is the progeny of some parental age groups. Every newborn has a probability *p*_*i*_ of coming from a parent of age *i*, such that ∑_*i*_*p*_*i*_ = 1. Note that our *N*_0_ is equivalent to Felsenstein’s (1971) *N*_1_, the number of individuals born every time unit. Thus, *N*_0_ is not an effective size, but a census size. The probability of a newborn to live to age *i* is given by *s*_*i*_ (*l*_*i*_ in Felsenstein, 1971), so that the number of individuals that survive to age *i* is given by *N*_*i*_ = *s*_*i*_*N*_0_.

For convenience, we firstly derive the equations for **Q** when generations overlap. We then calculate the equivalent **L** from which we derive equations to calculate the number of newborns *N*_0_. This, in turn, can be used to calculate *N*_*e*_, the effective population size per generation. We then describe how simulations were performed. Lastly, we analyse an available dataset of white crowned sparrows (*Zonotrichia leucophrys*).

### Analytical derivations for probabilities of joint IBD

The full derivation for the evolution of **Q** and **L** is given in Appendix A and Appendix B, respectively. For each of them, a full system of equations is derived, which includes the probabilities of joint IBD (without replacement for **Q** and with replacement for **L**). Such system initially includes probabilities of joint IBD across all cohorts. For instance, in the case of **Q**, it includes all *Q*_*ij*_’s, the probabilities of joint IBD between gametes of cohort *i* and cohort *j*. The system for **Q** reduces to

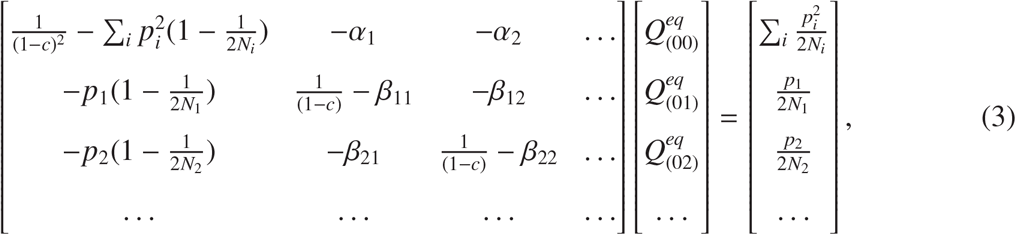

where β_*ik*_ = ∑_*j*_ δ_|*i*− *j*|,*k*_ *p*_*j*_ and α_*k*_ = ∑_*i*_ ∑_*j*_ δ_|*i*− *j*|,*k*_ *p*_*i*_ *p*_*j*_. The function δ_*i, j*_ is Kroenecker’s delta, that is, δ_*i,i*_ = 1 and δ_*i, j*_ = 0 for *i* ≠ *j*.

Similarly, as shown in detail in Appendix B, at equilibrium, **L** follows

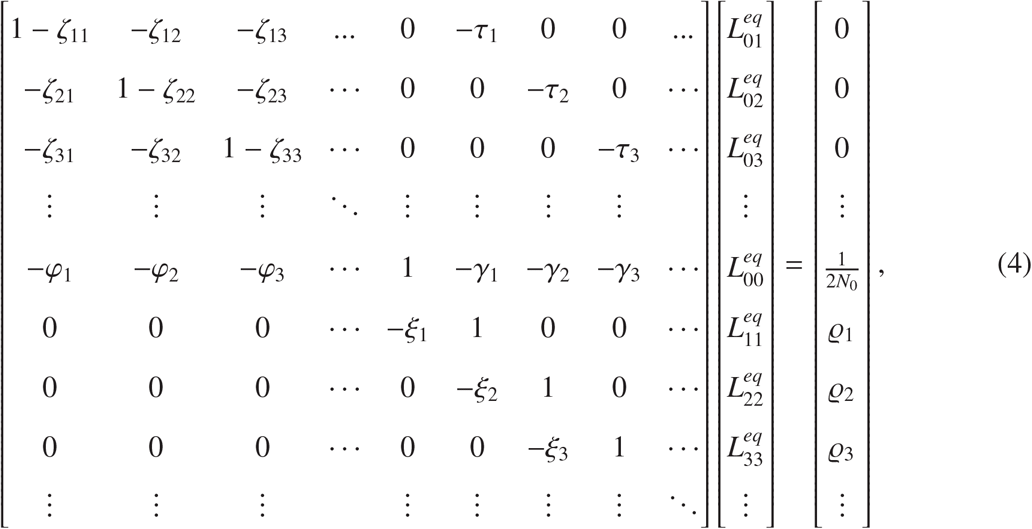

where ζ_*ik*_ = (1 − *c*) β_*ik*_, τ_*i*_ = (1 − *c*)*p*_*i*_, 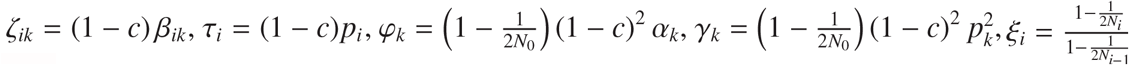 and 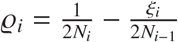.

### Estimation of N_0_ from linkage disequilibrium

As *E* (*r*^2^) = *L* (Sved & Feldman 1973), we can use the equations described in the previous section to estimate *N*_0_ from observed values of linkage disequilibrium. We thus use eq. 15, eq. 16, eq. 17 and eq. 18, but considering *N*_0_ as unknown and 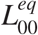 as observed for a given value of recombination frequency.

The equations for *N*_0_ are nonlinear but can be solved by iterating

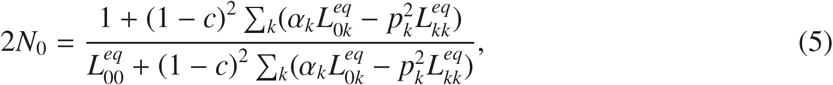

and the rest of identities in eq. (4), that is

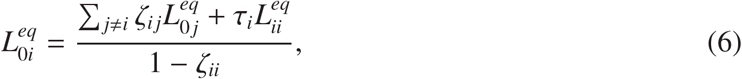

and

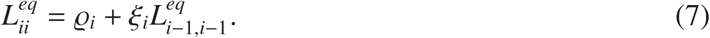

### Numerical analyses

We compared our analytical results with simulated results. For this purpose, we followed these steps in our simulations:

1. We drew the number of newborn gametes coming from each group age (*n*_*i*_) using multinomial deviates *n*_1_, *n*_2_, … ∼ *M* (*N*_0_, *p*_1_, *p*_2_, …), where *p*_*i*_ is the proportion of the 2*N*_0_ newborn gametes born from individuals from cohort *i*.
2. For each cohort *i* then
  a. We drew the number of recombined (*nrec*_*i*_) or non-recombined (*norec*_*i*_) gametes from a binomial deviate *nrec*_*i*_ ∼ *B* (*n*_*i*_, *c*).
  b. We drew the number of non-recombined progeny carrying each haplotype (*AB,ab,aB* and *Ab*) following a multinomial distribution *M* (*norec*_*i*_, *p*_*AB*_(*i*), *p*_*Ab*_(*i*), *p*_*aB*_(*i*), *p*_*ab*_(*i*)), where *p*_*AB*_(*i*),*p*_*Ab*_(*i*),*p*_*aB*_(*i*) and *p*_*ab*_(*i*) represent the haplotypic frequencies of cohort *i*.
  c. We drew the number of recombined progeny carrying each haplotype from a multinomial distribution *M* (*rec*_*i*_, *p*_*A*_(*i*)*P*_*B*_, *p*_*A*_(*i*)*P*_*b*_, *p*_*a*_(*i*)*P*_*B*_, *p*_*a*_(*i*)*P*_*b*_), where *p*_*A*_(*i*) and *p*_*a*_(*i*) are the allelic frequencies of cohort *i. P*_*B*_ and *P*_*b*_ represent the allelic frequencies of the whole population.
3. We calculated the haplotypic frequencies of newborn individuals by adding up gametes from steps 2b and 2c.
4. For each cohort *i*
  a. We then drew the number of individuals to be removed *died* ∼ *B* [*N*_*i*−1_, (*N*_*i*−1_ − *N*_*i*_) /*N*_*i*−1_] using *N*_*i*_ and *N*_*i*−1_,
  b. The number of individuals carrying each haplotype to be removed was then drawn from *M* (*died, p*_*AB*_(*i* − 1), *p*_*Ab*_(*i* − 1), *p*_*aB*_(*i* − 1), *p*_*ab*_(*i* − 1)), where *p* are the haplotypic frequencies at group age *i* − 1
  c. Individuals were eliminated to calculate the new haplotypic frequencies at each cohort *i*.

Recombined and non-recombined gametes at step 2a include gametes that have undergone an odd or even number of crossovers, respectively. These calculations were performed over 100,000 replicates, each of which with one pair of loci. Allelic frequencies were 1/2 at the beginning of each replicate.

### Simulations with many markers over several chromosomes

In order to take into account correlations between pairs of markers in the genome, and their effect on our estimates, we performed simulations for each scenario with either 1000 or 5000 markers distributed at random over 1, 5 or 10 chromosomes of 1M each. Allelic frequencies at the beginning of each replicate were drawn from a uniform distribution. For each newborn gamete, we drew two individuals from the existing cohorts, with probabilities following the contribution of each cohort (i.e., we drew a random number between 0 and 1, and if it falls between 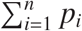 and 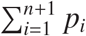 then the contributing parent comes from cohort *n* + 1). We generated recombinations within each of those individuals so that, on average, there was one recombination per chromosome. We repeated this process until *N*_0_ newborns were generated. After 500 generations, we calculated *r*^2^ for each segregating pair and using eqs.5, 6 and 7 we obtained an estimate of *N*_0_ for each pair. We then calculated the harmonic mean of *N*_0_ for each replicate.

### Application to real data

The programs required to infer *N*_0_ and *N*_*e*_ from observed *r*^2^ are available at https://github.com/angelesdecara/OverlapNeLD. We applied our method to available data from white-crowned sparrow (*Zonotrichia leucophrys nuttalli*) (Lipshutz *et al* 2017, 2016), whose age structure has been described in earlier works (Baker *et al* 1981; Waples & Yokota 2007). White crowned sparrows have approximately 5 fertile cohorts together with the newborns (i.e. 6 cohorts). Fertile individuals leave between 2.5 and 3.3 offspring per unit time, and the survival probability from newborns to one unit time old is 0.18, and then every unit time this probability is reduced by half roughly. These life-history values have been scaled such that population size remains constant. Data from 169 individuals are available for 6419 SNPs obtained using genotyping-by-sequencing (GBS). They can be classified in three different groups, depending on whether they are ssp. nuttalli, ssp. pugetensis or admixed. We calculated *N*_0_ and *N*_*e*_ using the values for the fraction of a cohort that survives to a certain age, and the mean number of offspring produced per generation by individuals of each age group from Waples & Yokota (2007), separately for each subspecies.

Linkage disequilibrium was estimated using the option `-geno-r2` of vcftools (Danecek *et al* 2011). Note that this measure of linkage disequilibrium is the correlation between allelic frequencies not taking into account gametic phases. Thus, it is an approximation of the gametic linkage disequilibrium. Furthermore, there is no chromosome scale reference genome for this species, and the number of data was small. We corrected this estimate of *r*^2^ to account for sample size following Weir & Hill (1980); Waples (2006); Waples & Do (2008), by the harmonic sample size for each pair of loci (see Waples & Do 2008, for further details).

As there is no available linkage map, we had no information on the positions of each SNP. Thus, we assumed that *c* = 0.5, that is, that all SNPs were independent. Thus, the results here obtained are likely to be downward biased. In order to infer the effective population size from *N*_0_, we followed the approach developed by Felsenstein (1971), which we solved with using a recursive formula. We compared our results with those obtained using Ne estimator (Waples & Do 2008) based on LD using all informative SNPs, equally assuming that SNPs are unlinked.

In order to provide a standard error, we performed a jackknife approach (Waples *et al* 2020; Jones *et al* 2016). Considering a sample size of *n* individuals, we selected *n* − 1 individuals *n* times. For each jackknife sample we calculated *R*^2^ (Waples & Do 2008), that is, the weighted correlation between genotype frequencies at different markers, corrected by sample size. Realistic sample sizes generate linkage disequilibrium due to sampling, and this needs to be corrected before applying any method to estimate *N*_*e*_ (Waples 2006; Waples & Do 2008). Once we have *R*^2^ for each jackknife sample, we obtained *N*_0_ and from there *N*_*e*_, which allowed us to obtain a standard error for both values. Similarly, we could have obtained the bounds of the confidence interval for *R*^2^ and use our equations to obtain the bounds for *N*_*e*_. Furthermore, we tested the effect of considering all loci, independently of their minor allele frequency (MAF), or only those with MAF greater than 0.05.

## Results

We compare here the results obtained analytically with those obtained computationally using multinomial distributions, following the above algorithm. The population here studied consists of three fertile cohorts or age groups, that produce *N*_0_ newborns. We studied the three following scenarios:

1. All contributions are the same (i.e. all *N*_*i*_’s take the same value of 1000 individuals) and 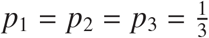.
2. All cohorts have the same number of individuals (all *N*_*i*_ = 1000 individuals), but they contribute differently to the number of newborns, due to different fertilities per age group, such that *p*_1_ = 0.3, *p*_2_ = 0.5 and *p*_3_ = 0.2.
3. Cohorts have different sizes (*N*_1_ = 1000, *N*_2_ = 800 and *N*_3_ = 500) and contribute differently *p*_1_ = 0.3, *p*_2_ = 0.5 and *p*_3_ = 0.2.

Fig. 3 shows analytical and simulated results for *L*_00_, in black and green respectively, for a population with 3 cohorts and *N*_0_ = 1000 individuals. The difference between the estimated (*L*_00_ as given by eqs. 15, 16 and 17) and the average *r*^2^ using 100,000 replicates (that is, estimated from *L*_00_ is larger as the contribution of one cohort increases as compared to the other cohorts, and furthermore, this difference increases with cohort age. In other words, the difference between estimated and observed when cohort 1 is the only one contributing is zero, but when cohort 3 is the most contributing this difference is larger than when cohort 2 is the most contributing.

As shown in tab. 1, we can see how linkage disequilibrium as measured by *r*^2^ is in very good agreement with the results obtained for *L*_00_ (eqs. 15, 16 and 17), which takes into account sampling with replacement, while *Q*_00_ (eq. 3) differs noticeably. This is the case for the three scenarios, with the difference between *r*^2^ and *Q*_00_ increasing with increasing complexity. Scenario 3 shows the largest differences between *r*^2^ and *L*_00_, but both measures are still in very good agreement, for a variety of recombination probabilities.

Estimates of *N*_0_ from *L*_00_ are shown in tab. 2, where we used the 100,000 results to provide the average, and then we used sets of either 100 or 500 pairs of loci to provide the standard deviation In both cases, the results are for *N*_0_ = 1000 individuals and we can see how the inferred *N*_0_ is mostly accurate within 15% deviation when 100 replicates are used and within 7% when 500 replicates are used.

Lastly, we have analysed the dependence on *c* and *N*_0_ (eq. 5). In fig. 4, we can see how the inferred *N*_0_ deviates from its true value of 1000 depending on the recombination fraction *c*. This deviation is stronger for tighter linkage, i.e. small *c*. In fig. 5, we can see how the inferred *N*_0_ has a high accuracy under all scenarios for all recombination rates.

A detailed analysis of the estimated *N*_0_ versus recombination frequency is shown in Fig. 4, where we can see the three scenarios studied. Overall, the larger the recombination frequency, the better the estimate of *N*_0_.

Similar results are shown in Fig. 5, where we can see the dependency of the estimated *N* with its true value for four recombination frequencies. The estimate provides a good result overall. Results on simulations with 1000 or 5000 markers randomly allocated in either 1, 5 or 10 chromosomes are shown in Table 3. The estimates are consistently good, and standard deviation decreases with number of chromosomes, as most measurements of *r*^2^ is between markers in different chromosomes, which results in the most accurate estimates of *N*_0_.

**Table 1:** Results for *r*^2^ evaluated for three scenarios, for different genetic distances (*d*) which correspond to different recombination rates (*c*). Results are given evaluated using multinomial distributions (“Empirical *r*^2^”), from *Q*_00_ or measured from *L*_00_. Scenario 1: All contributions are the same (i.e. all *N*_*i*_’s take the same value equal to 1000 individuals) and 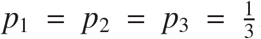. Scenario 2: All contributions are the same (all *N*_*i*_ = 1000 individuals), but *p*_1_ = 0.3, *p*_2_ = 0.5 and *p*_3_ = 0.2. Scenario 3: All contributions are different: *N*_1_ = 1000, *N*_2_ = 800, *N*_3_ = 500 and *p*_1_ = 0.3, *p*_2_ = 0.5 and *p*_3_ = 0.2. SD stands for standard deviation.

| Scenario | $d$ (cM) | $c$ | Empirical $r^2 \pm \text{SD} (\times 10^3)$ | $Q_{eq,00} \times 10^3$ | $L_{eq,00} \times 10^3$ |
| --- | --- | --- | --- | --- | --- |
| 1 | 5 | 0.0476 | $2.916 \pm 4.105$ | 2.406 | 2.905 |
| 1 | 10 | 0.0906 | $1.668 \pm 2.354$ | 1.166 | 1.666 |
| 1 | 20 | 0.1648 | $1.044 \pm 1.474$ | 0.551 | 1.051 |
| 1 | 50 | 0.3161 | $0.705 \pm 0.999$ | 0.201 | 0.701 |
| 1 | ∞ | 0.5 | $0.574 \pm 0.815$ | 0.071 | 0.571 |
| 2 | 5 | 0.0476 | $3.049 \pm 4.256$ | 2.539 | 3.038 |
| 2 | 10 | 0.0906 | $1.741 \pm 2.453$ | 1.234 | 1.733 |
| 2 | 20 | 0.1648 | $1.084 \pm 1.528$ | 0.586 | 1.086 |
| 2 | 50 | 0.3161 | $0.718 \pm 1.013$ | 0.216 | 0.715 |
| 2 | ∞ | 0.5 | $0.583 \pm 0.824$ | 0.078 | 0.578 |
| 3 | 5 | 0.0476 | $3.572 \pm 5.043$ | 2.940 | 3.439 |
| 3 | 10 | 0.0906 | $1.995 \pm 2.813$ | 1.435 | 1.935 |
| 3 | 20 | 0.1648 | $1.224 \pm 1.723$ | 0.687 | 1.187 |
| 3 | 50 | 0.3161 | $0.772 \pm 1.090$ | 0.257 | 0.757 |
| 3 | ∞ | 0.5 | $0.598 \pm 0.846$ | 0.094 | 0.594 |

**Table 2:** Results for estimated *N*_0_ evaluated for three scenarios, for different genetic distances (*d*) which correspond to different recombination rates (*c*). Results are given evaluated using *r*^2^ obtained via for *L*_00_, using 100 and 500 pair of loci. Scenario 1: All contributions are the same (i.e. all *N*_*i*_’s take the same value equal to 1000 individuals) and 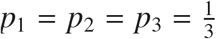. Scenario 2: All contributions are the same (all *N*_*i*_ = 1000 individuals), but *p*_1_ = 0.3, *p*_2_ = 0.5 and *p*_3_ = 0.2. Scenario 3: All contributions are different: *N*_1_ = 1000, *N*_2_ = 800, *N*_3_ = 500 and *p*_1_ = 0.3, *p*_2_ = 0.5 and *p*_3_ = 0.2.

| Scenario | $d$ | $c$ | Empirical $r^2 \times 10^3$ | Mean | SD<sub>100</sub> | SD<sub>500</sub> |
| --- | --- | --- | --- | --- | --- | --- |
| 1 | 5 | 0.0476 | 2.875 | 991.24 | 145.19 | 65.24 |
| 1 | 10 | 0.0906 | 1.657 | 996.31 | 143.85 | 63.40 |
| 1 | 20 | 0.1648 | 1.051 | 1005.66 | 145.45 | 63.97 |
| 1 | 50 | 0.3161 | 0.701 | 994.24 | 144.67 | 63.01 |
| 1 | ∞ | 0.5 | 0.574 | 996.16 | 140.77 | 65.58 |
| 2 | 5 | 0.0476 | 3.049 | 996.67 | 145.50 | 62.01 |
| 2 | 10 | 0.0906 | 1.733 | 993.18 | 139.80 | 58.52 |
| 2 | 20 | 0.1648 | 1.087 | 999.98 | 145.68 | 63.53 |
| 2 | 50 | 0.3161 | 0.713 | 995.94 | 152.03 | 63.74 |
| 2 | ∞ | 0.5 | 0.583 | 991.48 | 140.64 | 60.48 |
| 3 | 5 | 0.0476 | 3.540 | 956.48 | 141.35 | 60.35 |
| 3 | 10 | 0.0906 | 1.994 | 966.77 | 143.50 | 64.65 |
| 3 | 20 | 0.1648 | 1.227 | 968.07 | 138.63 | 60.84 |
| 3 | 50 | 0.3161 | 0.772 | 980.36 | 141.15 | 62.30 |
| 3 | ∞ | 0.5 | 0.598 | 992.85 | 138.76 | 62.15 |

**Table 3:** Mean *N*_0_ for each scenario using simulations of 1000 or 5000 markers in 1, 5 or 10 chromosomes. Standard deviation was calculated over 1000 replicates. Scenario 1: All contributions are the same (i.e. all *N*_*i*_’s take the same value equal to 1000 individuals) and 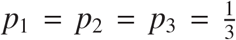. Scenario 2: All contributions are the same (all *N*_*i*_ = 1000 individuals), but *p*_1_ = 0.3, *p*_2_ = 0.5 and *p*_3_ = 0.2. Scenario 3: All contributions are different: *N*_1_ = 1000, *N*_2_ = 800, *N*_3_ = 500 and *p*_1_ = 0.3, *p*_2_ = 0.5 and *p*_3_ = 0.2.

| Scenario | Markers | Chromosomes | $N_0$ | SD |
| --- | --- | --- | --- | --- |
| 1 | 1000 | 1 | 999.86 | 15.88 |
| 1 | 5000 | 1 | 999.69 | 14.44 |
| 1 | 1000 | 5 | 999.70 | 5.34 |
| 1 | 5000 | 5 | 999.34 | 4.25 |
| 1 | 1000 | 10 | 999.46 | 4.25 |
| 1 | 5000 | 10 | 999.43 | 3.32 |
| 2 | 1000 | 1 | 998.77 | 15.99 |
| 2 | 5000 | 1 | 998.77 | 15.24 |
| 2 | 1000 | 5 | 999.38 | 5.63 |
| 2 | 5000 | 5 | 999.35 | 4.49 |
| 2 | 1000 | 10 | 999.69 | 4.83 |
| 2 | 5000 | 10 | 999.37 | 3.47 |
| 3 | 1000 | 1 | 999.46 | 17.48 |
| 3 | 5000 | 1 | 998.93 | 16.36 |
| 3 | 1000 | 5 | 998.98 | 6.12 |
| 3 | 5000 | 5 | 1000.00 | 4.91 |
| 3 | 1000 | 10 | 999.42 | 4.98 |
| 3 | 5000 | 10 | 999.76 | 3.90 |

*Application to real data: white crowned sparrows* Using the life-history parameters from Waples & Yokota (2007) and our exact equations, we inferred *N*_0_, and then, using the recursive equations from Felsenstein (1971), we used this result to infer *N*_*e*_. Results on inferred *N*_*e*_ are shown in Table 4. There we can see that the inferred *N*_*e*_ is rather low, which would indicate the need for a conservation programme. It is however possible that this low *N*_*e*_ is partly due to the assumption of independent SNPs, and using a different value of *c* would result in larger estimates of *N*_*e*_.

**Table 4:** Estimated *N*_0_ and *N*_*e*_ for white-crowned sparrow, together with Ne estimator results (Waples & Do 2008). *S* is the weighted harmonic mean of the sample size. *R*^2^ is a measure of linkage disequilibrium from unphased data, corrected by sample size.

| Individuals used | $S$ | Corrected $R^2$ | $N_0$ | $N_e$ | Ne estimator |
| --- | --- | --- | --- | --- | --- |
| Nuttalli | 33 | 0.000766 | $1418.88 \pm 94.73$ | $525.06 \pm 35.07$ | $433.4 \pm 28.96$ |
| Pugetensis | 71.1 | 0.000264 | $4117.43 \pm 116.63$ | $1524.01 \pm 43.17$ | $1260.6 \pm 35.57$ |

**Table 5:** Dependence on minor allele frequency (MAF) of the estimates of *N*_0_ and *N*_*e*_. A MAF *p* = 0 means no locus has been filtered, while at *p* = 0.05 those loci with alleles with minor allele frequency lower or equal to 0.05 have been removed from the analysis.

| Subspecies | $N_e$ | | Ne estimator | |
| --- | --- | --- | --- | --- |
| | $p = 0$ | $p = 0.05$ | $p = 0$ | $p = 0.05$ |
| Nuttalli | $525.06 \pm 35.07$ | $239.33 \pm 7.32$ | $433.4 \pm 28.96$ | $196.40 \pm 6.05$ |
| Pugetensis | $1524.01 \pm 43.17$ | $745.27 \pm 17.70$ | $1260.6 \pm 35.57$ | $615.51 \pm 14.62$ |

The estimates for *N*_*e*_ are considerably lower than those obtained by Lipshutz *et al* (2017) using fastsimcoal (Excoffier *et al* 2013), which uses the site frequency spectrum (SFS) to make demographic inferences. This discrepancy between estimates based on LD or those based on fastsimcoal using the SFS has been observed in other studies (Sovic *et al* 2019). The differences may be due to several factors, like the fact that Lipshutz *et al* (2017) assumed constant population sizes throughout long time-scales, and allowed for migration (Santiago *et al* 2020).

## Discussion

Following Sved (1971); Sved & Feldman (1973) and earlier work (Garcia-Cortes *et al* 2019), we have derived here exact equations for linkage disequilibrium in populations with overlapping generations. Combining this result with life-history parameters, we have inferred the number of newborns per generation, *N*_0_, which in turn, can be used to estimate effective population size, a key parameter in evolutionary and conservation studies. We have derived estimates per generation, not per year, and thus, we have not derived *N*_*b*_, the effective population size per year. The strength of our method is that we do not require any approximation, as it relies on an analytical approach. Our goal here was partly to take into account the developments made by Sved & Feldman (1973); Sved (2019) so that sampling with replacement is considered. Sved (2019) emphasised the linked identity by descent (*L*), which is conceptually equivalent to what he called earlier (Sved 1971) joint identity by descent (*Q*). Both identities are related by equations (13) and (14). Our goal was to develop a model-based method, and thus we have not compared our results with the heuristic approach of Waples *et al* (2014).

Our results show that using the linked identity by descent provides better estimates, which is in agreement with our earlier work (Garcia-Cortes *et al* 2019). We have further compared our exact results with those obtained via simulations, using multinomial distributions, and have obtained a very good agreement between them.

Our method is reliable and behaves well even with a strong age-structure, as seen on Fig. 4 and Fig. 5. Based on Fig. 4, the best estimates are obtained for loosely linked or unlinked markers. The good agreement can also be observed in Fig. 3, where we can see the good agreement between the estimated and the simulated size. It seems that the older the most contributing cohort, the largest the differences between the estimated and the simulated size. Even so, the estimates are very good and mostly within 10% relative error.

We have developed equations to infer *N*_*e*_ starting from *L*_00_, that is, linkage disequilibrium within newborns. Using such measure, we can derive all linkage disequilibria within and between cohorts, which leads to the inference of *N*_*e*_ by using the sets of equations(4). Alternatively, we could take a random sample from the population to estimate overall linkage disequilibrium. In that case, we could infer *N*_*e*_ combining this overall estimate of LD, which is the weighted sum of the linkage disequilibria between and within cohorts. This would add further complexity to the problem, as it would require equations(4) together with this weighted average.

Our method is based on a set of assumptions, and therefore, it has to be applied with caution. In order to derive analytical results, we have assumed constant population and cohort sizes, as well as constant life-history parameters. We have assumed random mating, and thus, variance in mean reproductive success is equal to the mean.

Finally, we have used our analytical results to infer *N*_*e*_ in the white-crowned sparrow, for which genetic and demographic data were available (Baker *et al* 1981; Lipshutz *et al* 2017). The results shown in Tab. 4 are in line with our previous results. We had shown for non-overlapping generations that the estimator of Weir & Hill (1980) was downward biased (Garcia-Cortes *et al* 2019). In Table 4 we observe a similar trend, where the results obtained with Ne estimator, based on the estimator developed by Weir & Hill (1980), are smaller than those obtained with our method. We have to emphasise that this dataset suffers from a small sample size, lack of linkage map and using an estimate of *r*^2^, as gametic phases are unavailable. Furthermore, we have used life-history parameters that were estimated in one population of the subspecies. Our results are limited by the information we have, and thus should be taken with caution.

Obiously, our results are still limited by the assumptions here made, like constant age structure and constant population size. We warn researchers working in species or populations who are aware of huge expansions or bottlenecks or fluctuating size that this is probably not the most appropiate method to analyse their data. Most importantly, life-history parameters need to be adjusted so that population size is constant (Waples & Yokota 2007). Our method does not work and will give incorrect results if this premise is not held. Some populations may have extremely succesful individuals who leave a large number of offspring. This is a well-known issue when estimating effective population size. We have assumed a Poisson distribution in the reproductive success, and thus, deviations from this distribution may bias the result.

Genomic data are now easily accessible, yet, there are two issues which may make our method not straightforward. The first one is lack of life-history parameters, which require long-term studies to characterise survival and fertility. The second one is the lack of reference genomes. Our method can be applied to any type of genomic data, and ideally a recombination map should be available. Otherwise, we recommend filtering the data by minor allele frequencies (Jones *et al* 2016, and references therein), as rare alleles are known to bias the inference, and assume free recombination for unmapped markers from RADseq or GBS. For those cases where scaffolds are available, keep one SNP from each scaffold, and depending on genome size and number of markers and scaffolds, analyse them as unlinked. Based on our simulations, we think that there where sequence data are available, we believe using about 10000 good quality evenly sparsed throughout the genome is more than enough.

This study shows the potential of our method to make estimates of effective population size in wild species using the available programs at http://github.com/angelesdecara/OverlapNeLD.

## Acknowledgements

This work was funded by Ministerio de Economía y Competitividad grant number AGL2016-75942-R, by the LabEx BCDIV (ANR-10-LABX-0003-BCDiv), within the framework of the program “Investing for the future” (ANR-11-IDEX-0004-02) and by the Agence Nationale de la Recherche project AGRHUM (ANR-14-CE02-0003). We are most grateful to Elizabeth Derryberry, Sara Lipshutz, Amy Luo and Ryan Waples for feedback on the data and the manuscript. The cluster Finis Terrae II at Centro de Computación de Galicia was used for this work.

## Appendix A: Dynamics of Q throughout generations

Following the notation for *Q* of Sved (1971), we denote *Q*_*ij*_ as the probability of joint homozygosity between a pair of gametes from age groups *i* and *j* and *Q*_*ii*_ is such probability between two different gametes of the same age group *i, i*.*e*. two gametes sampled without replacement. Then, at a given time interval,

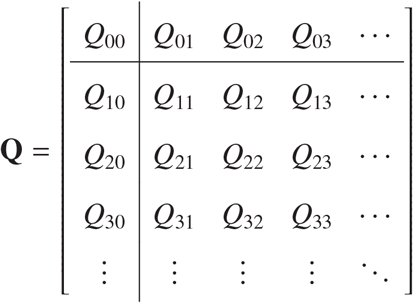

where 0 is the group of newborns. The definition of matrix **Q** here mimics the definition of the heterozygosity matrix in Felsenstein (1971).

We denote 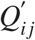 as the elements of **Q** at the next time unit. Every time unit the *Q* at each cohort can be written in terms of contributions from other *Q*s at other cohorts, and analogously for between cohorts *Q*_*i j*_. In fig. 1, we show the five possible origins of a pair of random gametes in a population with two parental age groups. This can be easily extended for (*n*^2^ + 3*n*)/2 contributions with *n* parental groups. The recursive formula to obtain 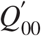, i.e. *Q*_00_ at the next time unit, given the elements of **Q** can be obtained adding all contributions on fig. 1. This can be generalised to any number of age groups such that

**Figure 1:**
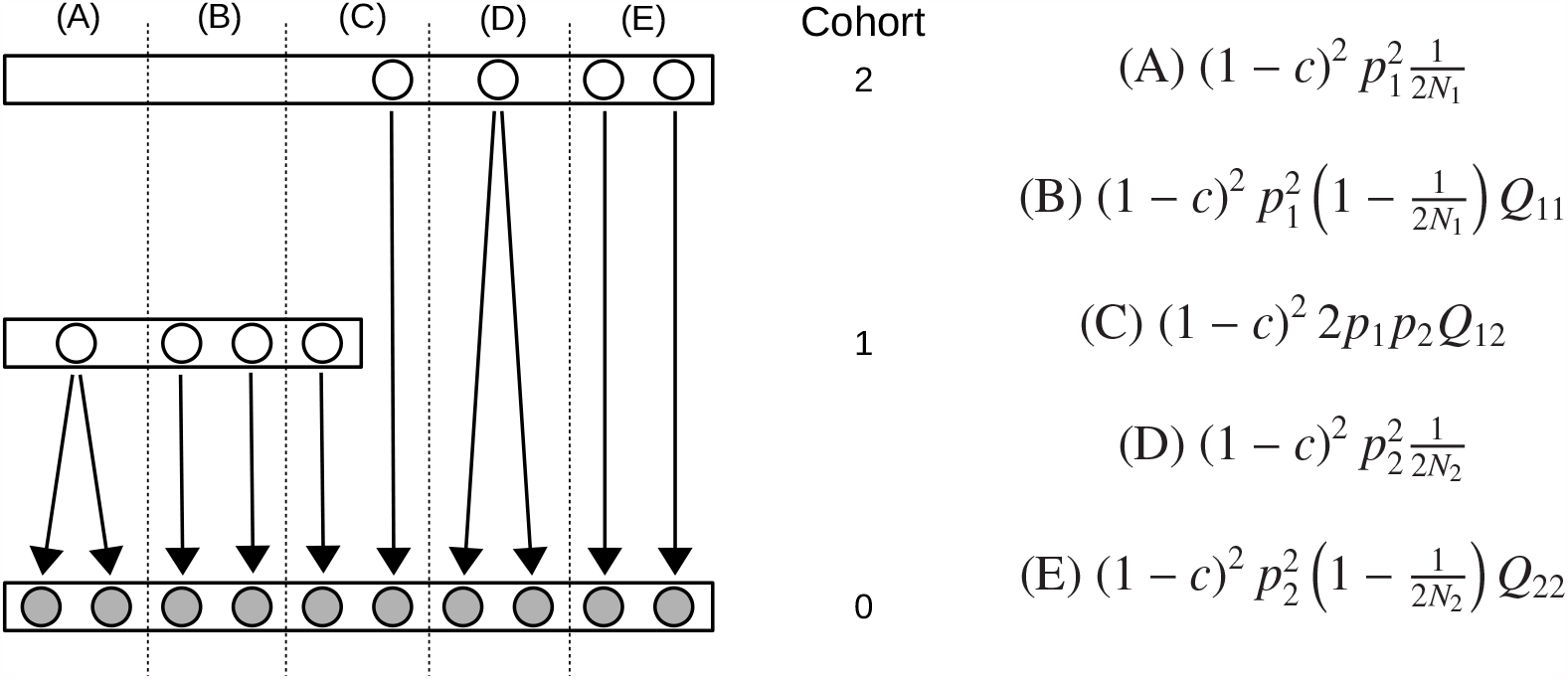
Inheritance of gametes in a scenario with two cohorts. Left: (A) Probability of two gametes (on gray) to be offspring of the same gamete one cohort back in time. (B) Probability of two gametes to be offspring from two independent gametes one cohort back in time Case (C) shows two gametes that are offspring from gametes from two different cohorts. In case (D) two gametes are offspring from the same gamete two cohorts back in time. Lastly, in case (E), two gametes who descend from two different gametes two cohorts back in time. On the right, the probabilities of each case (A)-(E).

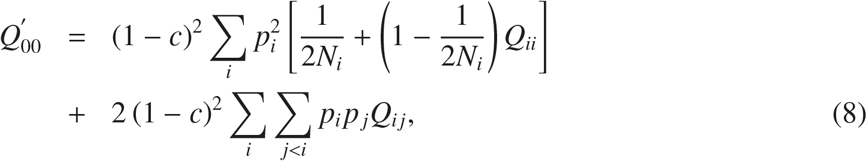

where the term 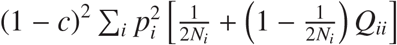 corresponds to the sum of the contribution of cases (a), (b), (d) and (e) in fig. 1 while the term 2 (1 − *c*)^2^ ∑_*i*_ ∑_*j*<*i*_ *p*_*i*_ *p*_*j*_*Q*_*i j*_ corresponds to case (c) extended for an arbitrary number of cohorts. In order to derive the probabilities of each case, and following Sved (1971), the term (1 − *c*)^2^ accounts for no crossover in either maternal or paternal meiosis, while 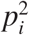 or *p*_*i*_ *p*_*j*_ is the probability of a pair of gametes to come from the same or different age groups. Lastly, 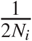 and 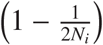 denote the probabilities that ancestral gametes to be identical or not, respectively.

Fig. 2 shows the contributions of the three possible cases for both 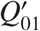 and 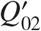 (or *n* + 1 contributions for *n* age groups). Adding all contributions for an arbitrary number of age groups, the recursive formula to obtain each 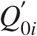 is

**Figure 2:**
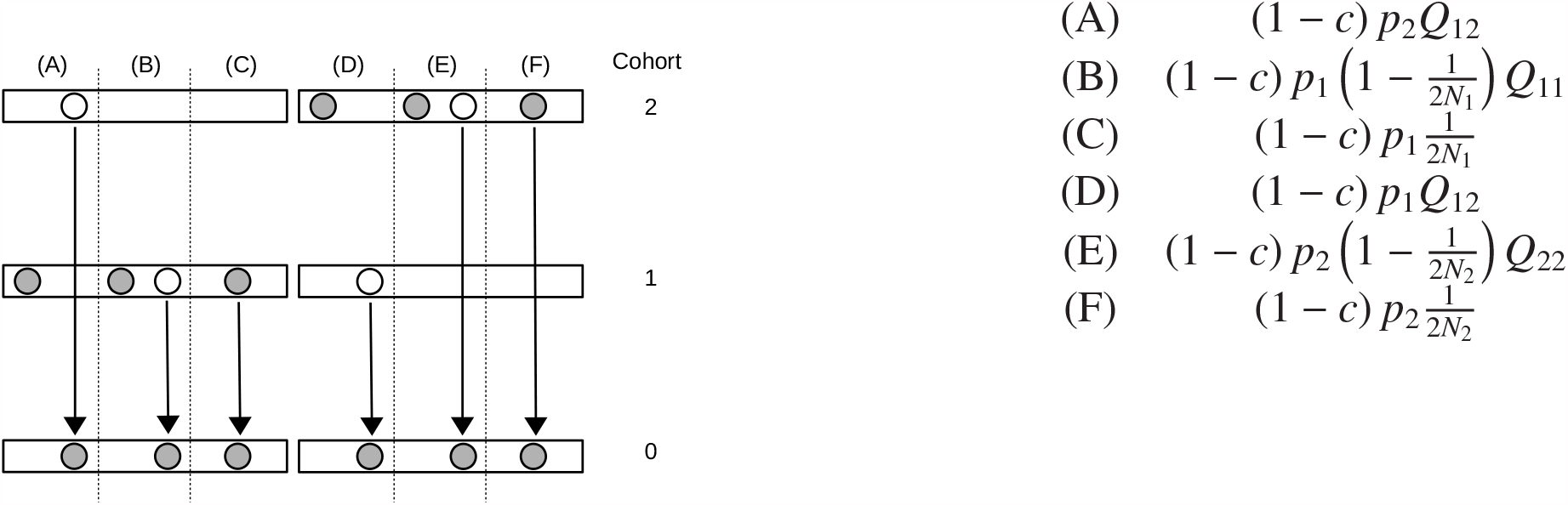
Probability of two gray gametes to be IBD given that one is offspring from the white gamete. Cases A to C show all conditional IBD between two gametes of successive cohorts, while cases D to F illustrate conditional IBD between two gametes born two cohorts apart. Probabilities are shown on the right.

**Figure 3:**
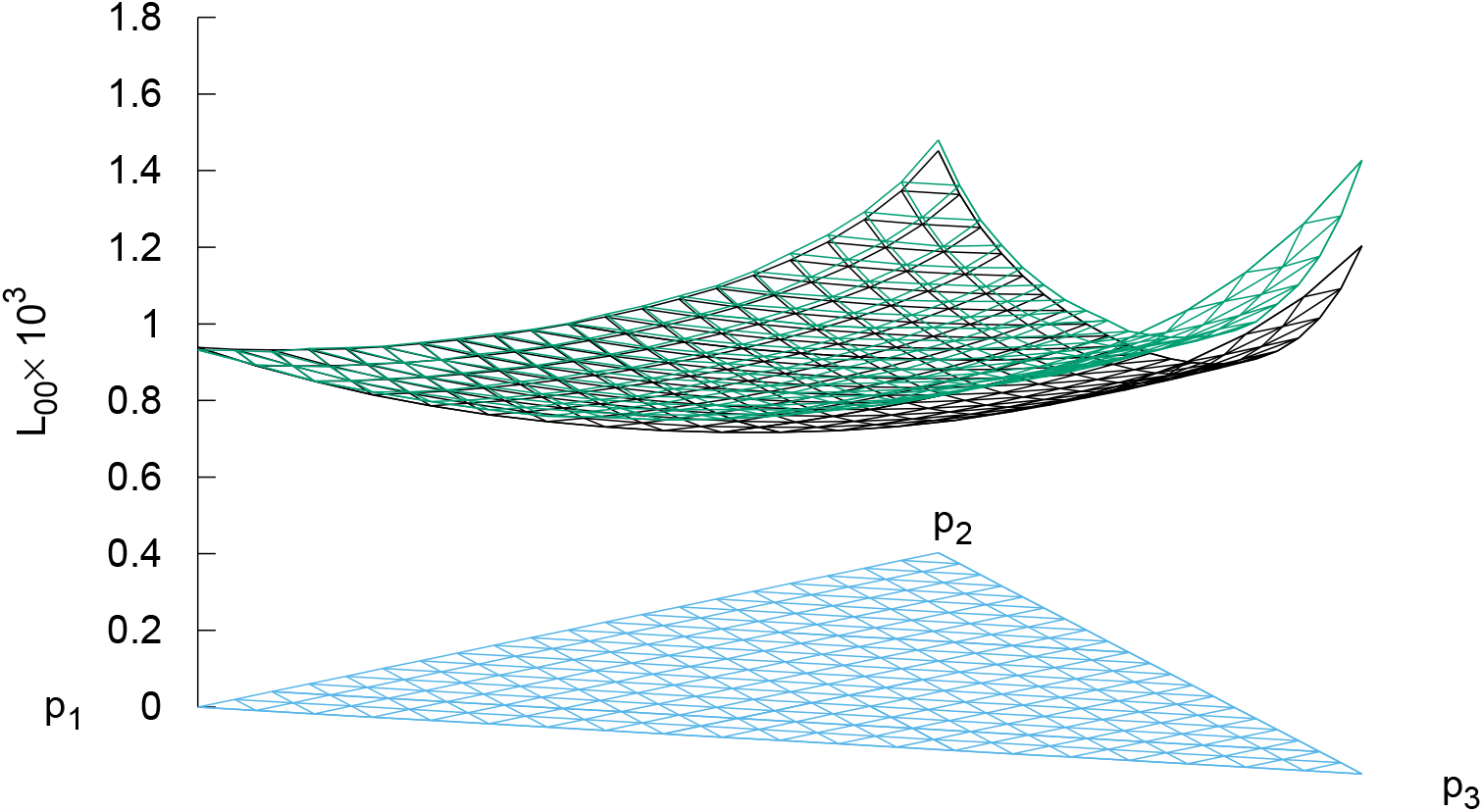
Equilibrium values for *L*_00_ obtained via analytical results (black), simulations using multinomial distributions (green). *N*_0_ = 1000 individuals and loci are assumed to be at 50cM. The triangle axis represent the contribution from each cohort *p*_*i*_ to the pool of newborns, which satisfy 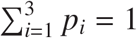.

**Figure 4:**
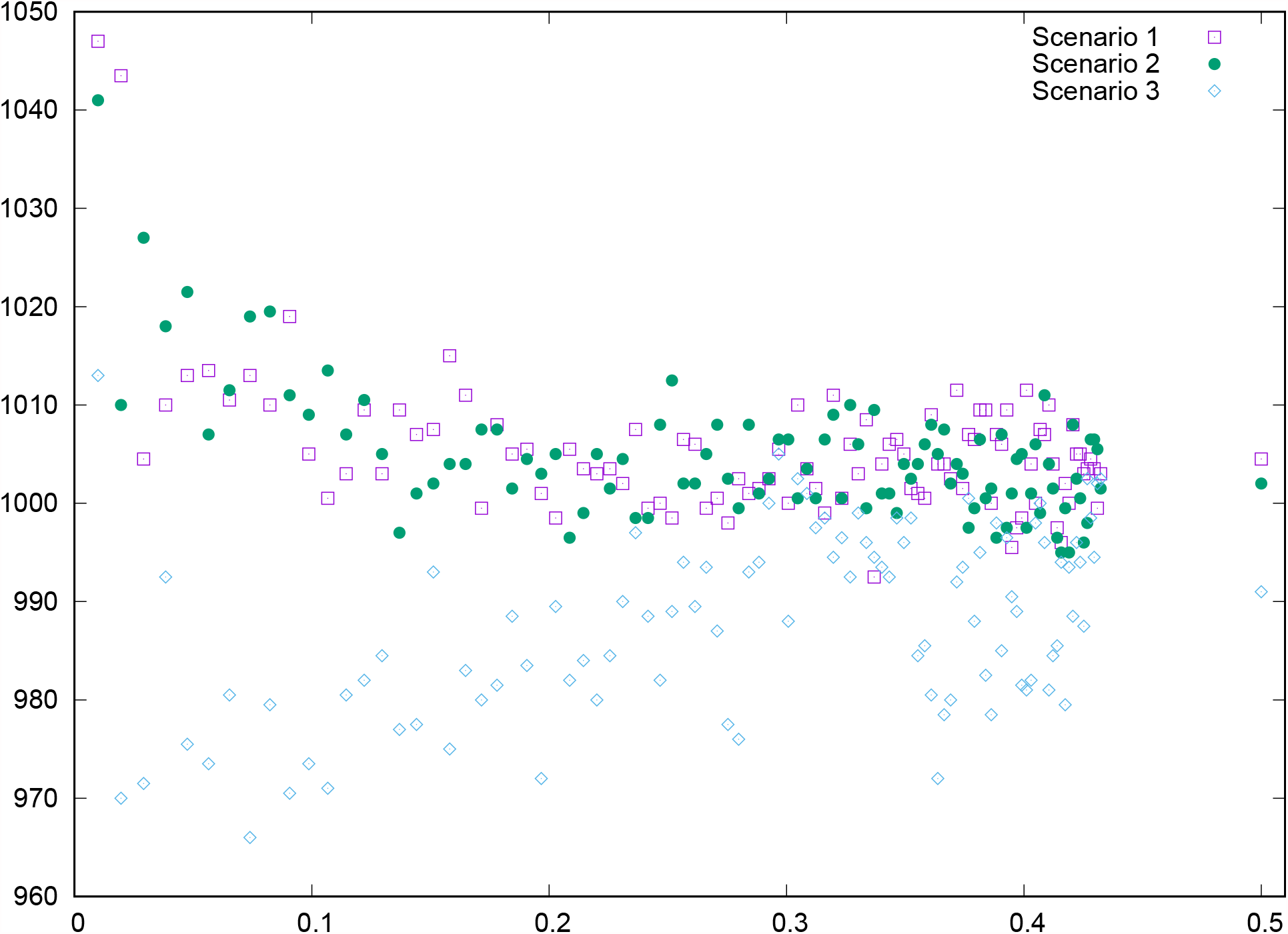
Inferred *N*_0_ for a population with true *N*_0_ = 1000 individuals, under three different scenarios. Values are averages over 100000 replicates.

**Figure 5:**
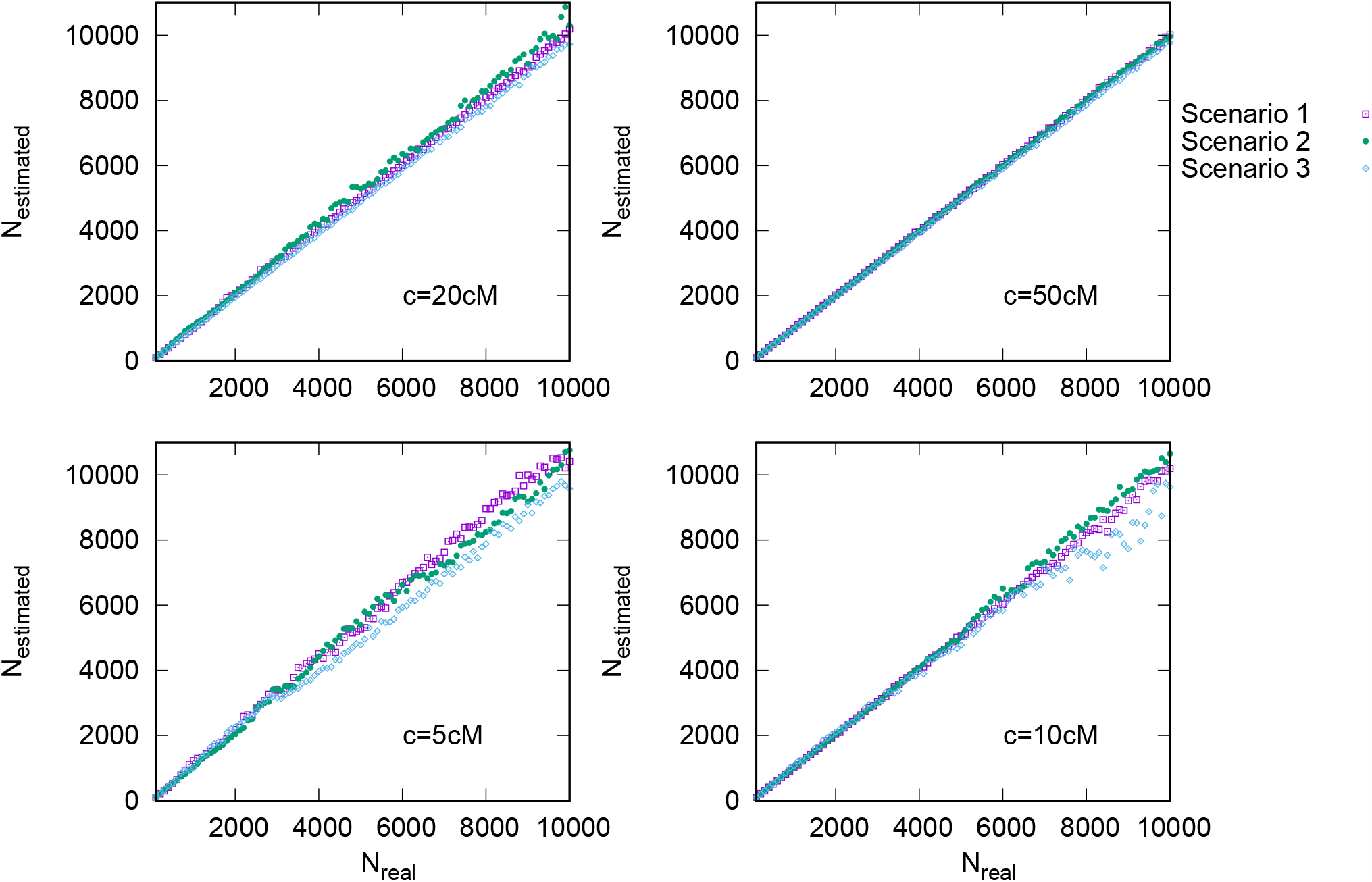
Inferred *N*_0_ for a population with varying *N*_0_ individuals, under three different scenarios. Recombination distances are fixed at 5cM, 10 cM, 20 cM and 50 cM, from bottom to top. Values are averages over 100000 replicates.

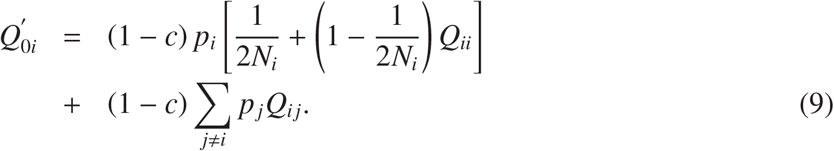

Given that individuals from cohort *i* will move onto cohort *i* + 1 in the next time unit, the joint homozygosity between survivors of cohort *i* and cohort *j* in a time unit remains the same in the next time unit. Therefore, the remaining elements in **Q**^′^ are

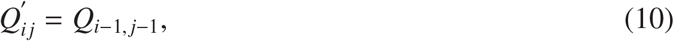

and

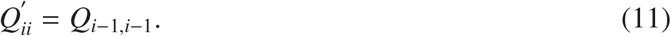

The recursive formulae given by eq. (8) to eq. (11) reach an equilibrium **Q**^*eq*^ where drift and recombination balance out. At equilibrium 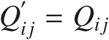 and, considering eq. (10) and eq. (11), it follows that *Q*_00_ = *Q*_11_ = *Q*_22_, *Q*_01_ = *Q*_12_ = *Q*_23_, *Q*_02_ = *Q*_13_ and so on. Thus, the matrix given by

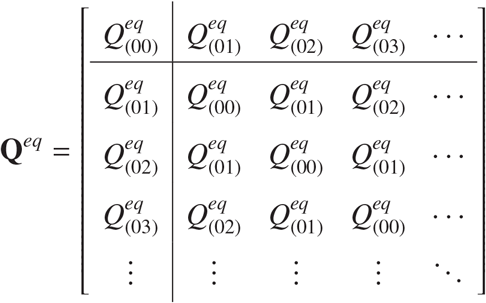

and equations (8) and (9) can be solved using

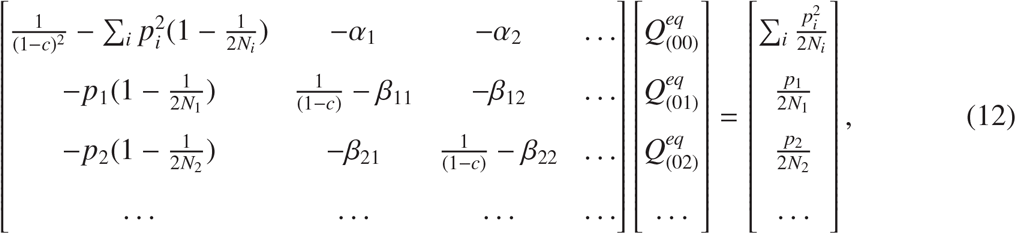

where β_*ik*_ = ∑_*j*_ δ_|*i*− *j*|,*k*_ *p*_*j*_ and α_*k*_ = ∑_*i*_∑_*j*_ δ_|*i*− *j*|,*k*_ *p*_*i*_*p*_*j*_. The function δ_*i, j*_ is Kroenecker’s delta, that is, δ_*i,i*_ = 1 and δ_*i, j*_ = 0 for *i* ≠ *j*.

## Appendix B

Dynamics of L throughout generations

Here, we derive the equations for *L*, the probability of linked identity by descent, at two positions, as defined by Sved & Feldman (1973) and Sved (2019). We define *L*_*i j*_ as the LIBD of a pair of gametes from age groups *i* and *j* and *L*_*ii*_ as the LIBD of two gametes of age group *i*, drawn at random with replacement. Then, at any given generation, *L* is given by

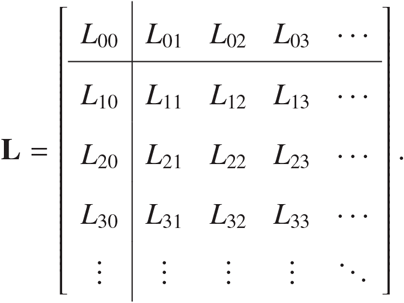

Following the relationships derived in appendix **C**, the relationship between *Q* and *L* is thus:

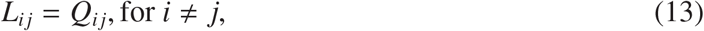

and

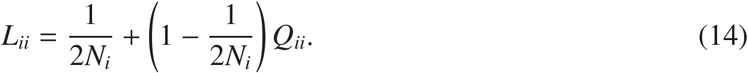

To calculate 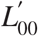, we replace eq. (13) and eq. (14) in eq. (8):

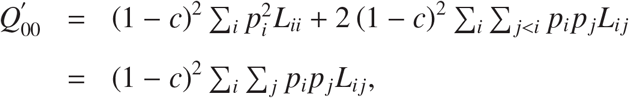

and replacing in eq. (14), we obtain that

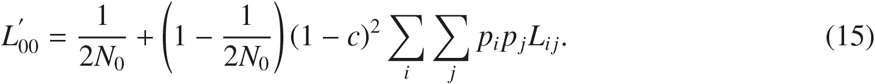

Similarly, to calculate 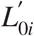, we replace both eq. (13) and eq. (14) in eq. (9), obtaining

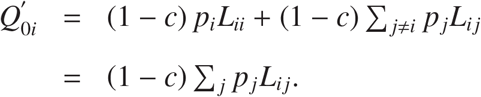

We can use these results in eq. (13):

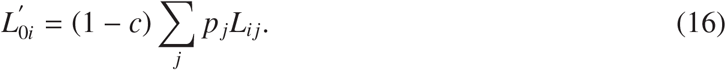

It then follows from eq. (10) and eq. (13) that

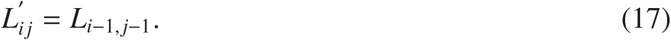

In fact, sampling with or without replacement are equivalent when considering gametes from different age groups. Finally, to calculate 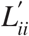 we rewrite eq. (14) as

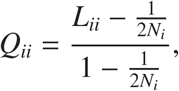

and replace it in eq. (11), which results in

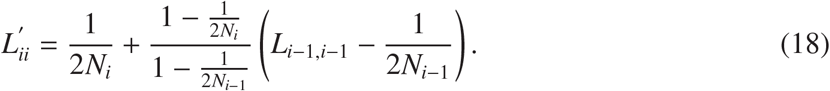

Using all the above results, eq. (18) can be rewritten as

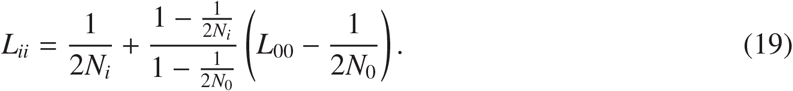

At equilibrium, 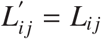 and, following eq. (17), *L*_01_ = *L*_12_ = *L*_23_, *L*_02_ = *L*_13_, … and so on. Thus,

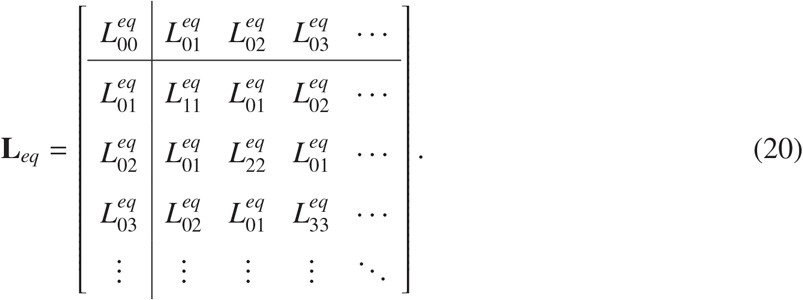

Therefore, at equilibrium

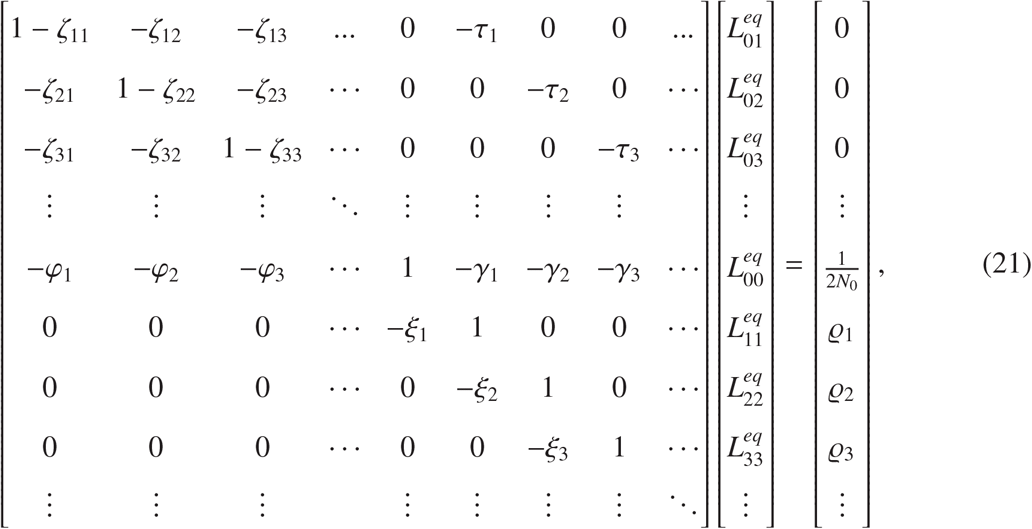

where ζ_*ik*_ = (1 − *c*) β_*ik*_, τ_*i*_ = (1 − *c*)*p*_*i*_, 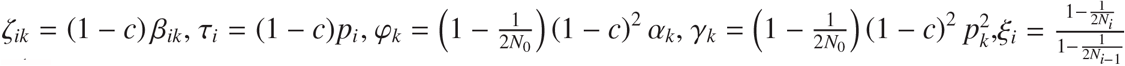 and 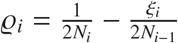.

It is well known that sampling with replacement usually leads to equations simpler than those derived considering sampling without replacement. For this reason, eq. (15) and eq. (16) are simpler than eq. (8) and eq. (9). Nevertheless, the number of equations to solve for *L*^*eq*^ is larger than the number of equations required to solve *Q*^*eq*^, which accounts for differences between the diagonal elements of matrix (20).

## Appendix C: Note on relation between Sved (1971) and Sved & Feldman (1973)

Both methods are equivalent, but the first develops equations for *Q* while the latter iterates *L*. As it seems *L* = *r*^2^, both methods are equivalent if

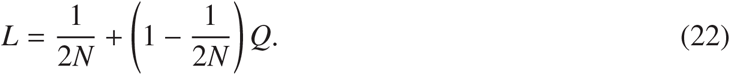

Sved (1971) develops the following recursive equation

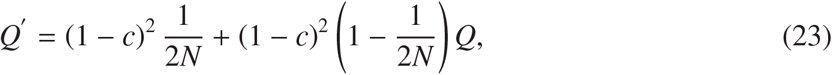

while Sved & Feldman (1973) develop recursive equations for

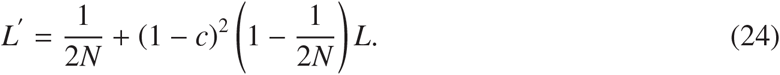

We can show this by replacing *L* and *L*^′^ in eq. (24) using eq. (22):

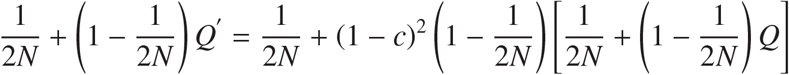

and, replacing *Q*^′^, it coincides with eq. (23):

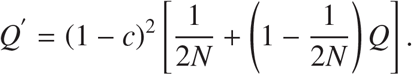

